# AMPKα2 is a skeletal muscle stem cell intrinsic regulator of myonuclear accretion

**DOI:** 10.1101/2022.11.02.514556

**Authors:** A Kneppers, M Theret, S Ben Larbi, L Gsaier, A Saugues, C Dabadie, A Ferry, K Sakamoto, R Mounier

## Abstract

Due to the post-mitotic nature of skeletal muscle fibers, adult muscle maintenance relies on dedicated muscle stem cells (MuSCs). In most physiological contexts, MuSCs support myofiber homeostasis by contributing to myonuclear accretion, which requires a coordination of cell-type specific events between the myofiber and MuSCs. Here, we addressed the role of the kinase AMPKα2 in the coordination of these events supporting myonuclear accretion. We demonstrate that AMPKα2 deletion impairs skeletal muscle regeneration. Through *in vitro* assessments of MuSC myogenic fate and EdU-based cell tracing, we reveal a MuSC-specific role of AMPKα2 in the regulation of myonuclear accretion, which is mediated by phosphorylation of the non-metabolic substrate BAIAP2. Similar cell tracing *in vivo* shows that AMPKα2 knockout mice have a lower rate of myonuclear accretion during regeneration, and that MuSC-specific AMPKα2 deletion decreases myonuclear accretion in response to myofiber contraction. Together, this demonstrates that AMPKα2 is a MuSC-intrinsic regulator of myonuclear accretion.

## INTRODUCTION

Skeletal muscle tissue constitutes approximately 45% of the total body weight in healthy humans, is a major determinant of the basal metabolic rate, and has well-recognized endocrine functions^1,2^. A low skeletal muscle mass is associated with metabolic disorders such as insulin resistance and type 2 diabetes, and is a risk factor for cardiovascular diseases and mortality^3–5^, illustrating the importance of skeletal muscle for whole body homeostasis. In addition, skeletal muscle quality and quantity are vital prerequisites for breathing, locomotion and performing daily tasks^6^. These functions are supported by the excitability and contractility of the myofibers, which are syncytia composed of hundreds of post-mitotic nuclei. Due to the post-mitotic nature of myofibers, adult muscle maintenance relies on dedicated skeletal muscle stem cells (MuSCs; a.k.a. satellite cells), which through their exit from quiescence, expansion, differentiation, and subsequent fusion, contribute to *de novo* myofiber formation after injury^7^.

Most commonly used models to study MuSC function invoke myofiber death by intramuscular injection with myotoxic agents, or by physical procedures such as freeze injury^8^, leading to a subsequent *de novo* myofiber formation. However, in humans complete *de novo* myofiber formation is exceedingly rare, since myofiber damage is usually contraction-induced and can lead to segmental myofiber necrosis^9,10^. As such, the recovery from myofiber damage in humans does not just rely on the fusion among MuSCs, but also relies on the fusion of MuSCs with pre-existing myofibers – *a process called ‘myonuclear accretion’*. In addition to its role in the resolution of myofiber injury, myonuclear accretion was recently demonstrated to specifically contribute to adaptive remodelling in response to physical exercise^11,12^. Moreover, myonuclear accretion occurs continuously during life to support skeletal muscle homeostasis in both the young and adult organism^13^. Thus, while common experimental models assess fusion among MuSCs during *de novo* myofiber formation, in most physiological contexts MuSCs support skeletal muscle homeostasis by their contribution to myonuclear accretion.

MuSC fusion is a tightly regulated process, which is illustrated by the array of proteins that play a critical role^14^, and the absolute requirement of muscle specific fusogens Myomaker (MYMK) and Myomixer (MYMX) to overcome the forces that prevent spontaneous membrane fusion^15–18^. Many of the components of the fusion machinery have been identified, but the physiological regulators of MuSC fusion remain largely unknown. Perhaps unsurprisingly, recent evidence points towards a distinct regulation of MuSC-MuSC fusion and myonuclear accretion^e.g.,19–21^. For example, Eigler *et al*. proposed that Ca^2+^/Calmodulin-dependent Protein Kinase II (CaMKII) activation in growing myotubes specifically facilitates myonuclear accretion^21^. In addition, Serum Response Factor (SRF)/ Cyclooxygenase-2 were shown to promote myonuclear accretion through the transcriptional regulation of the MuSC recruitment factor Interleukin-4 (IL-4) in myofibers^20^. These studies illustrate that myonuclear accretion requires a coordination of cell-type specific events between the myofiber and MuSCs. However, it remains unknown if there is a central molecular regulator that coordinates these events.

We postulate that such a molecular regulator may be contraction-induced, as physical exercise is a robust physiological trigger of myonuclear accretion. Skeletal muscle contraction imposes an acute demand on myofiber Ca^2+^ fluxes and energy turnover. Interestingly, several pathways that distinctly regulate myonuclear accretion converge on the key energy sensor and master regulator of cell metabolism ‘5’-AMP-activated protein kinase’ (AMPK). Indeed, AMPK is known to be regulated by Ca^2+^ signalling via Ca^2+^/Calmodulin-dependent Protein Kinase Kinase 2 (CaMKK2). Furthermore, AMPK is a transcriptional and posttranscriptional regulator of Peroxisome Proliferative Activated Receptor, γ, Coactivator 1 α (PGC1α)^22^, which has recently been suggested to participate in the regulation of myonuclear accretion upon endurance type training^23^.

AMPK is a heterotrimer composed of a catalytic subunit (α) and two regulatory subunits (β and γ)^22^. The predominant AMPKα isoform in skeletal muscle is AMPKα2, which is also more robustly activated upon physical exercise^24^. We therefore wanted to address whether AMPKα2 has a role in the coordination of cell-type specific events supporting myonuclear accretion during the re-establishment of skeletal muscle homeostasis.

In uninjured, resting adult skeletal muscle, AMPKα2 is predominantly expressed in the myofibers. We demonstrate that AMPKα2 expression is progressively increased during MuSC-mediated myogenesis, and that MuSC-specific AMPKα2 deletion impairs skeletal muscle regeneration. Through *in vitro* assessments of the MuSC myogenic fate, we show that AMPKα2 knockout specifically decreases MuSC-myofiber fusion (*i*.*e*., myonuclear accretion). Furthermore, using 5 ethynyl 2’ deoxyuridine (EdU)-based cell tracing *in vitro*, we reveal a MuSC-specific role of AMPKα2 in the regulation of myonuclear accretion, which is mediated by the phosphorylation of the non-metabolic substrate BAR/IMD domain containing adaptor protein 2 (BAIAP2; a.k.a. IRSp53). Similar EdU-based cell tracing *in vivo* shows that AMPKα2 knockout mice have a lower rate of myonuclear accretion during regeneration, and that MuSC-specific AMPKα2 deletion decreases the relative rate of myonuclear accretion in response to myofiber contractions induced by neuromuscular electrical stimulation (NMES). Together, our analyses demonstrate that AMPKα2 is a MuSC-intrinsic regulator of myonuclear accretion.

## RESULTS

### AMPKα2 deletion impairs skeletal muscle regeneration

Using the integrated single-cell and single-nucleus transcriptomic database scMuscle^25^, we observed that AMPKα1 (*Prkaa1*) is expressed in various cell types present in the skeletal muscle, while AMPKα2 gene (*Prkaa2*) expression is largely confined to myonuclei (Figure S1A). Similarly, we have previously shown that AMPKα1 activity was present, while AMPKα2 activity is absent in quiescent MuSCs^26^. Interestingly, during *in vitro* myogenesis of FACS-sorted wildtype MuSCs, *Prkaa2* expression was progressively increased (Figures S1B). To assess AMPK activity during skeletal muscle regeneration *in vivo, Tibialis Anterior* (TA) muscles of wildtype mice were subjected to Cardiotoxin (CTX)-induced injury (Figure 1A). Paralleling muscle cell-specific *in vitro* assessments, AMPKα2 activity was nearly absent at 2 days post injury (d.p.i.) when myofibers underwent a complete degeneration, and was gradually restored during myofiber regeneration at 7 and 14 d.p.i. (Figure 1B). Conversely, AMPKα1 activity was increased after 4 d.p.i., coinciding with MuSC proliferation and immune cell infiltration (Figure 1C).

**Figure 1.**
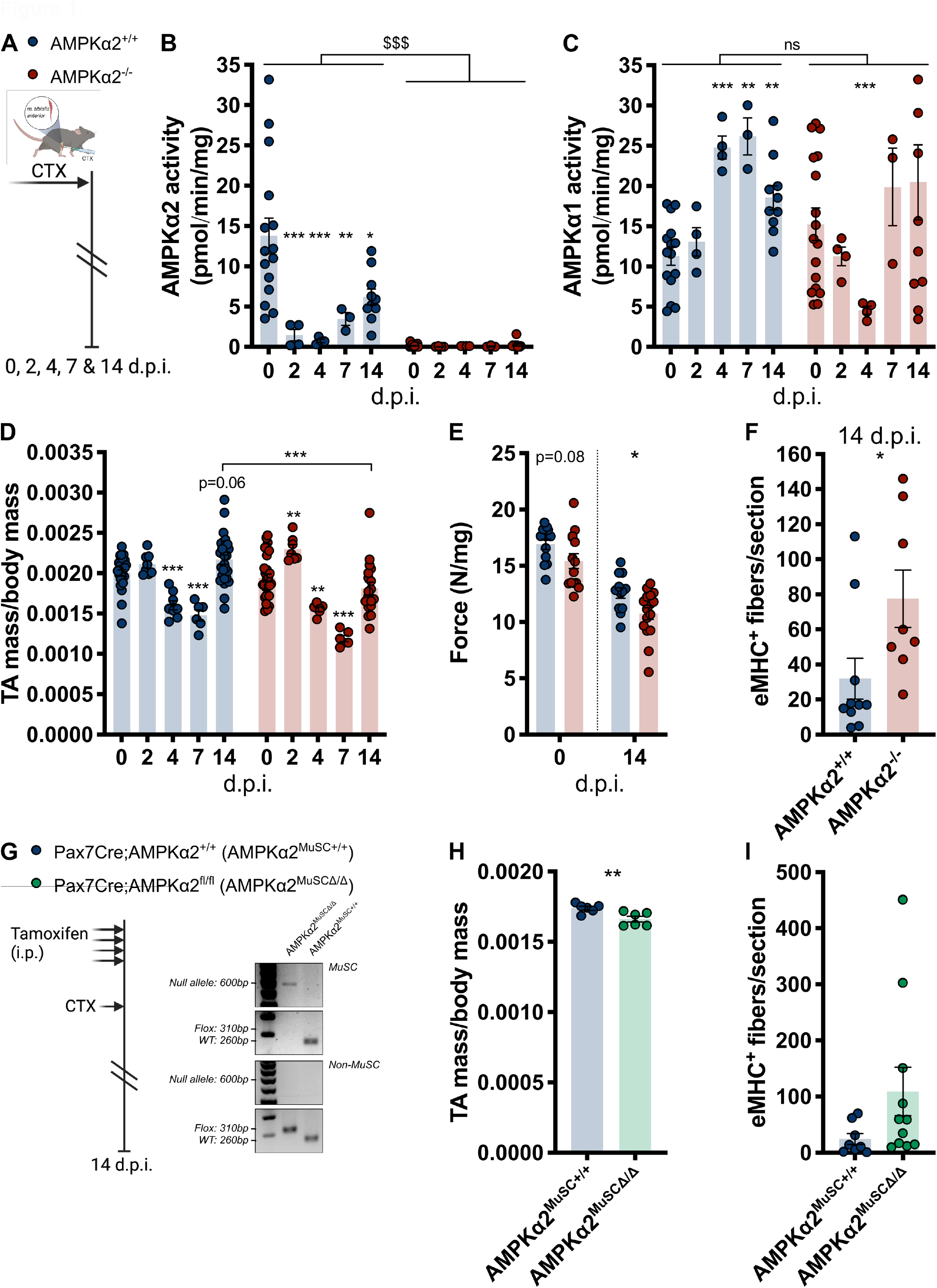
Role of AMPKα2 in skeletal muscle regeneration. Skeletal muscle regeneration in whole body AMPKα2^-/-^ mice (A-E). A. Schematic of Cardiotoxin (CTX)-induced skeletal muscle regeneration model and analysis endpoints at indicated days post injury (d.p.i.). B. Relative AMPKα2 activity during skeletal muscle regeneration. C. Relative AMPKα1 activity during skeletal muscle regeneration. D. Relative *Tibialis Anterior* (TA) mass during skeletal muscle regeneration. E. Relative *in situ* TA maximal force production before and 14 d.p.i. F. Number of eMHC stained fibers per section at 14 d.p.i. Skeletal muscle regeneration in mice after MuSC-specific AMPKα2 deletion (F-H). G. Schematic of Tamoxifen-induced AMPKα2 deletion and subsequent CTX-based skeletal muscle regeneration model (left panel), and verification of MuSC-specific recombination in FACS-sorted MuSC (CD45/CD31/Sca1^-^;CD34/α7int^+^) *versus* non-MuSC (CD45/CD31/Sca1^+^) extracted from hindlimb muscles of Pax7Cre;AMPKα2^+/+^ (AMPKα2^+/+^) and Pax7Cre;AMPKα2^fl/fl^ (AMPKα2^Δ/Δ^) mice at day 0 (*i*.*e*., 3 weeks after the first tamoxifen injection) (right panel). H. Relative TA mass at 14 d.p.i. I. Number of eMHC stained fibers per section at 14 d.p.i. Bars represent mean +/- SEM. Two-way ANOVA; ^$$$^p<0.001 genotype effect. Two-tailed, unpaired t-test; *p<0.05, **p<0.01, ***p<0.001, compared to 0 d.p.i., to AMPKα2^+/+^ control, or between indicated bars. See also Figure S1.

To study the role of AMPKα2 in skeletal muscle regeneration, mice lacking functional AMPKα2 (AMPKα2^-/-^) were utilized. As anticipated, AMPKα2 activity was not detectable throughout the regenerative process (Figure 1B). Importantly, AMPKα1 activity did not display a compensatory increase in AMPKα2^-/-^ TA (Figure 1C). Compared to control wildtype mice, AMPKα2^-/-^ TA was unaffected before injury, suggesting that AMPKa2 is not required for muscle development (Figure 1D). However, AMPKα2^-/-^ muscle had a lower relative mass and *in situ* force production at 14 d.p.i. (Figures 1D and 1E), while muscle fatigue remained unaffected (Figures S1C and S1D). Moreover, AMPKα2^-/-^ muscle contained a greater number of fibers expressing the regeneration marker embryonic Myosin Heavy Chain (eMHC) at 14 d.p.i., which returned to normal by 28 d.p.i., indicating an impairment in regeneration (Figures 1F and S1E). Yet, the number of Paired Box 7 (Pax7)^+^ cells per fiber and the number of fibers per section were unaltered before injury, and at 28 d.p.i. (Figures S1F and S1G), demonstrating that myofiber re-formation and MuSC maintenance were unaffected in AMPKα2^-/-^ TA.

To assess if the role of AMPKα2 during skeletal muscle regeneration is muscle cell-intrinsic, Pax7CreER^T2/+^;AMPKα2^fl/fl^ mice were treated with tamoxifen to induce AMPKα2 deletion in MuSCs and their progeny (AMPKα2^MuSCΔ/Δ^), and then subjected to CTX-induced injury (Figures 1G and S1H). AMPKα2^MuSCΔ/Δ^ mice had a lower relative TA mass, and a higher number of eMHC^+^ fibers at 14 d.p.i. (Figures 1H and 1I), indicating that AMPKα2 plays a muscle cell-intrinsic role during skeletal muscle regeneration.

### AMPKα2 is a regulator of myoblast fusion

To dissect the muscle cell-intrinsic role of AMPKα2, myogenesis was modeled *in vitro* using FACS-sorted MuSCs (Figure S2A). EdU incorporation during the first 48 hours of culture was unaffected while EdU incorporation into MuSC progeny (*i*.*e*., myoblasts) was lower in AMPKα2^-/-^ cultures (Figures S2B and S2C), suggesting that AMPKα2 does not regulate MuSC activation but modulates the subsequent rate of proliferation. In parallel, myoblasts were induced to differentiate and fuse to form *in vitro* generated myofibers (*i*.*e*., myotubes) for up to 48 hours, leading to a progressive increase in the fusion index of both AMPKα2^+/+^ and AMPKα2^-/-^ cultures (Figure 2A). However, at 48 hours of *in vitro* myogenesis, the fusion index remained lower in AMPKα2^-/-^ cultures (Figure 2A). Conversely, treatment with a specific allosteric AMPK activator 991^27,28^ increased the fusion index (Figure 2B), confirming a muscle cell-intrinsic role of AMPKα2 during myogenesis *in vitro*.

**Figure 2.**
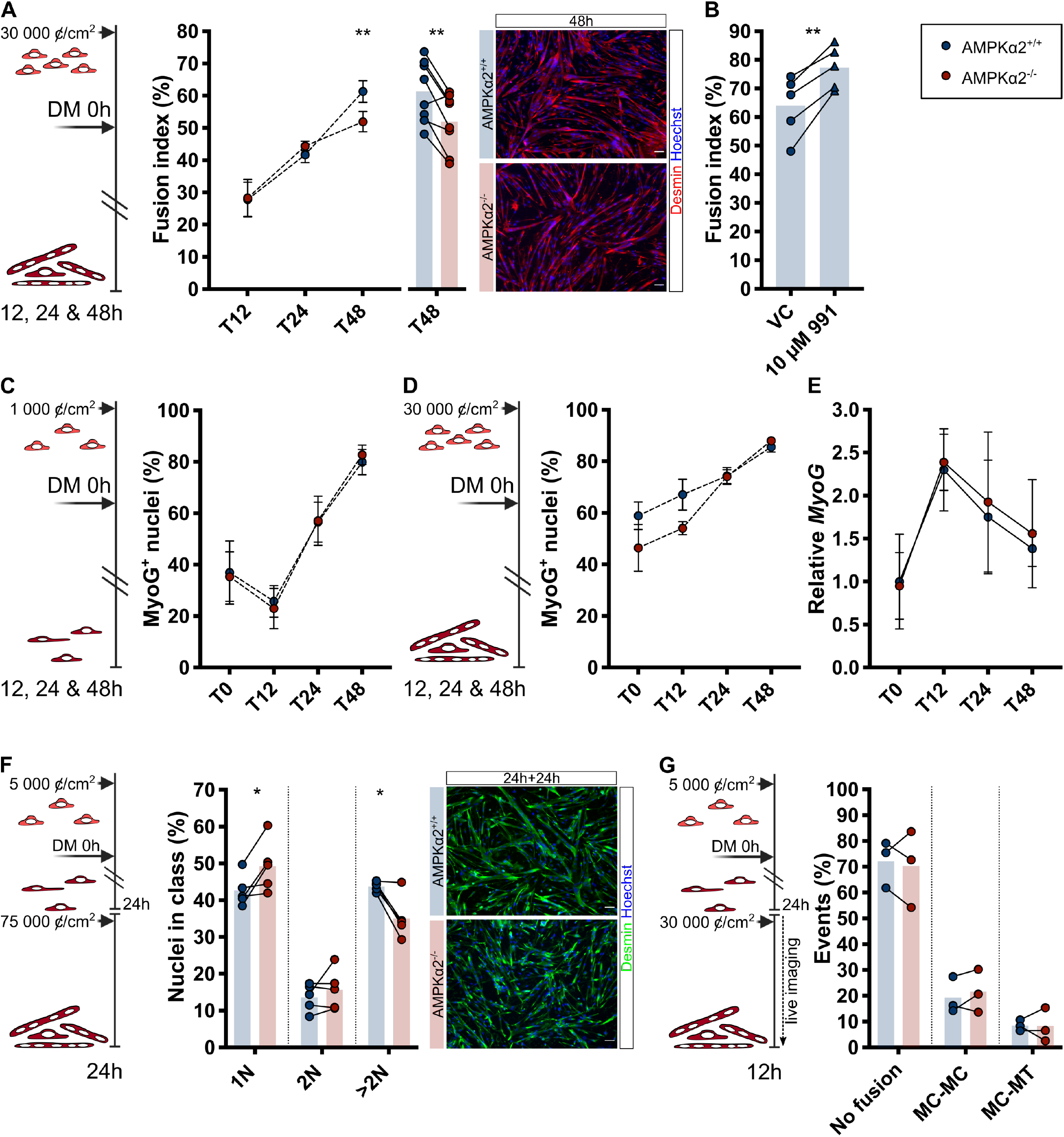
Role of AMPKα2 in the regulation of myoblast fusion *in vitro*. Left panels show schematic representation of experiments. A. Fusion index at indicated times of *in vitro* myogenesis, with insert of paired representation of the data at 48 hours (T48, middle panel), and representative images (right panel). B. Fusion index after 48 hours of *in vitro* myogenesis in cultures treated with 10 µM of the AMPK activator 991 or vehicle control (VC). C. Percentage of Myogenin (MyoG) stained nuclei at indicated times of differentiation in non-fusing cultures. D. Percentage MyoG stained nuclei at indicated times during *in vitro* myogenesis. E. Relative *MyoG* gene expression during *in vitro* myogenesis. F. Percentage of nuclei in cells with 1 nucleus (1N), 2 nuclei (2N) or >2 nuclei (>2N) after 24 hours of fusion of pre-differentiated cells (middle panel), and representative images (right panel). G. Fusion events scored during 12 hours of life imaging of fusing cultures of pre-differentiated cells. MC=myocytes, MT=myotubes. Bars represent mean. In line chart, dots represent mean +/- SEM. Two-tailed, paired t-test, *p<0.05, **p<0.01, compared to AMPKα2^+/+^ control, or to VC. Length of white scale bars represent 50 µm. See also Figure S2.

Metabolic alterations are known key drivers of MuSC differentiation^29^. Strikingly, the absence of AMPKα2 – which is known to act as a metabolic sensor and coordinator of metabolic processes – did not affect the muscle cell-intrinsic differentiation capacity, as the number of unfused cells expressing Myogenin (MyoG) increased similarly in AMPKα2^+/+^ and AMPKα2^-/-^ cultures (Figure 2C). Moreover, the percentage of MyoG^+^ nuclei and *MyoG* gene expression were unaffected in AMPKα2^-/-^ cultures during *in vitro* myogenesis (Figures 2D and 2E). Given the impaired fusion index despite unaffected differentiation in AMPKα2^-/-^ cultures, we sought to assess the muscle cell-intrinsic fusion capacity of AMPKα2^-/-^ cells. To this end, we differentiated myoblasts at low density to obtain differentiated mono-nucleated cells (*i*.*e*., myocytes), and subsequently cultured them at high density to isolate the process of fusion, as performed previously^30^. This assay resulted in a higher percentage of unfused cells (1N), and a lower fusion index in AMPKα2^-/-^ cultures due to a reduced presence of nuclei in cells with >2 nuclei (Figure 2F). Interestingly, the occurrence of cells containing 2 nuclei – resulting from the fusion between two mononucleated cells – was unaffected in these cultures (Figure 2F), and in cultures subjected to the previously described protocol for *in vitro* myogenesis (Figure S2D). To directly assess the fusion between two mononucleated cells, fusion events were scored during 12 hours live imaging of high-density cultured myocytes. We observed no difference in the rate of fusion between two mononucleated cells in AMPKα2^-/-^ cultures (Figure 2G, Video S1), which suggests that the core fusion machinery is unaffected. Indeed, gene expression of the fusogens *Mymk* and *Mymx* were unaffected in AMPKα2^-/-^ cultures (Figures S2E and S2F). Moreover, the motility parameters ‘velocity’ and ‘directness’, which influence the migration step of the fusion process, were unaltered (Figure S2G and Video S2). Together, this shows that AMPKα2 is a regulator of myocyte fusion, specifically controlling the formation of multinucleated myotubes.

### AMPKα2 is a myocyte-intrinsic regulator of myonuclear accretion

The general paradigm is that during skeletal muscle development and regeneration, the initial establishment of myofibers by MuSC-MuSC fusion is followed by myonuclear accretion to support the formation of multinucleated cells^31–33^. To directly address the role of AMPKα2 in the regulation of myonuclear accretion *in vitro*, we performed co-culturing between myotubes (MT) and myocytes (MC) (Figure 3A, Video S3.1-4). Myocyte-derived nuclei were traced by EdU incorporation at the myoblast stage, before initiation of co-culturing. After 24 hours of incubation, EdU incorporation was similar between AMPKα2^+/+^ and AMPKα2^-/-^ myocytes (Figure 3B). However, co-culture of AMPKα2^-/-^ myocytes with AMPKα2^+/+^ myotubes resulted in a lower myonuclear accretion percentage and a higher percentage of unfused AMPKα2^-/-^ myocytes (Figures 3C and 3D). Moreover, treatment with the AMPK activator 991 increased the myonuclear accretion percentage in co-cultures with AMPKα2^+/+^ myocytes, while this pro-fusion effect was absent in co-cultures with AMPKα2^-/-^ myocytes (Figures 3E and 3F). Conversely, when myocytes were co-cultured with AMPKα2^-/-^ myotubes the myonuclear accretion percentage and the percentage of unfused myocytes were unaffected (Figures S3A and S3B). In addition, the myonuclear accretion response to 991 treatment in co-cultures with AMPKα2^-/-^ myotubes was similar to that in co-cultures with AMPKα2^+/+^ myotubes (Figures S3C, S3D, 3E and 3F). Together, this shows that AMPKα2 is a myocyte-intrinsic regulator of myonuclear accretion.

**Figure 3.**
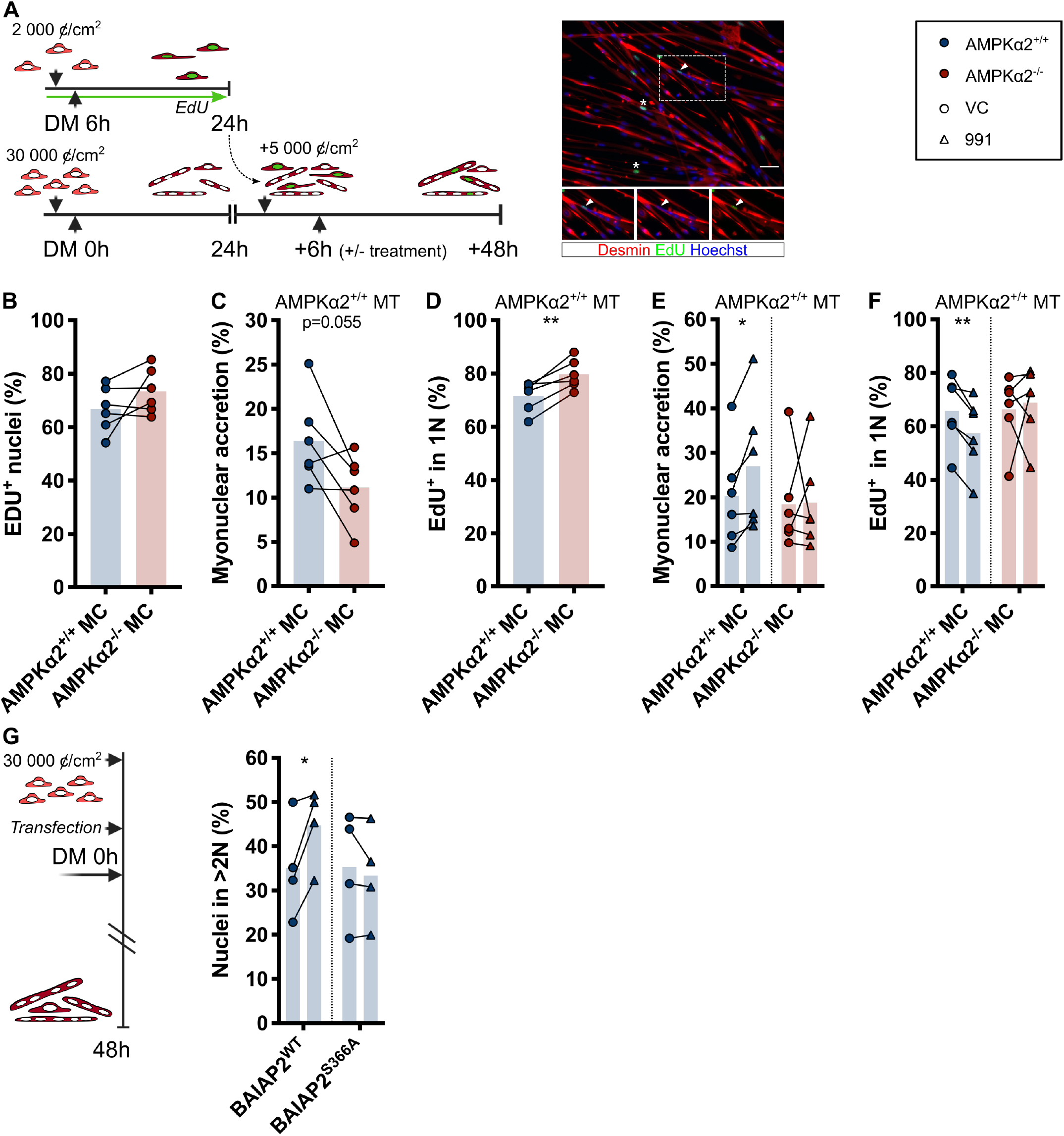
Myocyte-intrinsic regulation of *in vitro* myonuclear accretion by AMPKα2. A. Schematic of co-culturing assay between 24 hours differentiated myotubes and EdU-labeled pre-differentiated myocytes (left panel), and representative image (right panel). Arrow head indicates contribution of EdU^+^ myocytes to myonuclear accretion, stars indicate mononucleated EdU^+^ myocytes. Length of white scale bar represents 50 µm. MC=myocytes, MT=myotubes. B. Percentage EdU stained myocytes before start of co-cultures. Co-cultures between AMPKα2^+/+^ myotubes and AMPKα2^+/+^ *versus* AMPKα2^-/-^ myocytes (C-F). C. Percentage contribution of EdU^+^ myocytes to myonuclear accretion after 48 hours of co-culturing. D. Percentage EdU^+^ myocytes that remain mononucleated after 48 hours of co-culturing. E. Percentage contribution of EdU^+^ myocytes to myonuclear accretion after 48 hours of co-culturing with 10 µM of the AMPK activator 991 or vehicle control (VC). F. Percentage EdU^+^ myocytes that remain mononucleated after 48 hours of co-culturing with 10 µM 991 or vehicle control (VC). G. Percentage of nuclei in cells with >2 nuclei (>2N) after 48 hours of *in vitro* myogenesis of myoblasts transfected with BAIAP2 WT or BAIAP2^S366A^. Bars represent mean. Two-tailed, paired t-test, *p<0.05, **p<0.01, compared to AMPKα2^+/+^ control, or to VC. See also Figure S3.

Chemical genetic screens have identified BAIAP2 as a novel target of AMPKα2^34,35^. An *in vitro* study using the mouse myoblast cell line C2C12 previously identified BAIAP2 as a negative regulator of myogenesis^36^. Moreover, a GWAS associated *BAIAP2* gene variants to weight loss in patients with chronic obstructive pulmonary disease (COPD)^37^ – a disease linked with whole body and skeletal muscle wasting^38^. AMPKα2 directly phosphorylates BAIAP2 at Ser366^34,35^, which can regulate its cellular localization^39^. Moreover, regulation of the EPS8-BAIAP2 complex localization was recently linked to the progression of mammalian cell fusion^40^. We therefore explored the role of BAIAP2 in AMPKα2-mediated regulation of myonuclear accretion. *Baiap2* gene expression was strongly suppressed during *in vitro* myogenesis in both AMPKα2^+/+^ and AMPKα2^-/-^ cells (Figure S3E). However, transfection with the phospho-deficient mutant form BAIAP2^S366A^ prevented the pro-fusion effect induced by 991 (Figure 3G), suggesting that BAIAP2 mediates the regulation of fusion by AMPKα2.

### AMPKα2 is a MuSC-intrinsic regulator of myonuclear accretion in response to myofiber contractions induced by NMES

After CTX injury, MuSCs contribute to myofiber regeneration by MuSC-MuSC fusion and subsequent myonuclear accretion. To distinguish the MuSC contribution to these events *in vivo*, proliferating cells were traced by EdU labeling during the early (−1–5 d.p.i.) *versus* later phase of regeneration (5–14 d.p.i.) (Figure S4A). The timing of EdU labeling did not affect myonuclear number or localization (Figure S4B). Importantly, EdU labeling from 5-14 d.p.i. resulted in ∼20% of EdU^+^ myonuclei, that predominantly localize at the periphery of the myofibers (Figures S4C and S4D), strongly suggesting their contribution through myonuclear accretion^41^.

Using EdU labeling from 5-14 d.p.i., we observed a lower number of EdU^+^ nuclei co-stained with the myofiber nucleus marker Pericentriolar Material 1 (PCM1) in AMPKα2^-/-^ mice than in AMPKα2^+/+^ controls at 14 d.p.i., indicating a lower rate of myonuclear accretion *in vivo* in the absence of AMPKα2 (Figures 4A and 4B). In contrast, we observed no difference in the number of EdU^-^ myofiber nuclei in AMPKα2^-/-^ mice *versus* controls, showing that the first phase of regeneration including MuSC-MuSC fusion is unaffected *in vivo* (Figure 4B). Importantly, the total number of EdU^+^ nuclei below the basal lamina was reduced in AMPKα2^-/-^ mice (Figure S4E), confirming the lower rate of MuSC proliferation observed *in vitro* (Figure S2C). To explore the contribution of this proliferation defect to alterations in the myonuclear accretion rate, we expressed the EdU^+^/PCM1^+^ nuclei as a percentage of the total of EdU^+^ nuclei below the basal lamina. The resulting myonuclear accretion percentage is high (∼70%) and unaffected in AMPKα2^-/-^ mice (Figure 4C), suggesting that the myonuclear accretion defect in 14 days regenerated muscle is primarily caused by impaired MuSC proliferation in AMPKα2^-/-^ mice.

**Figure 4.**
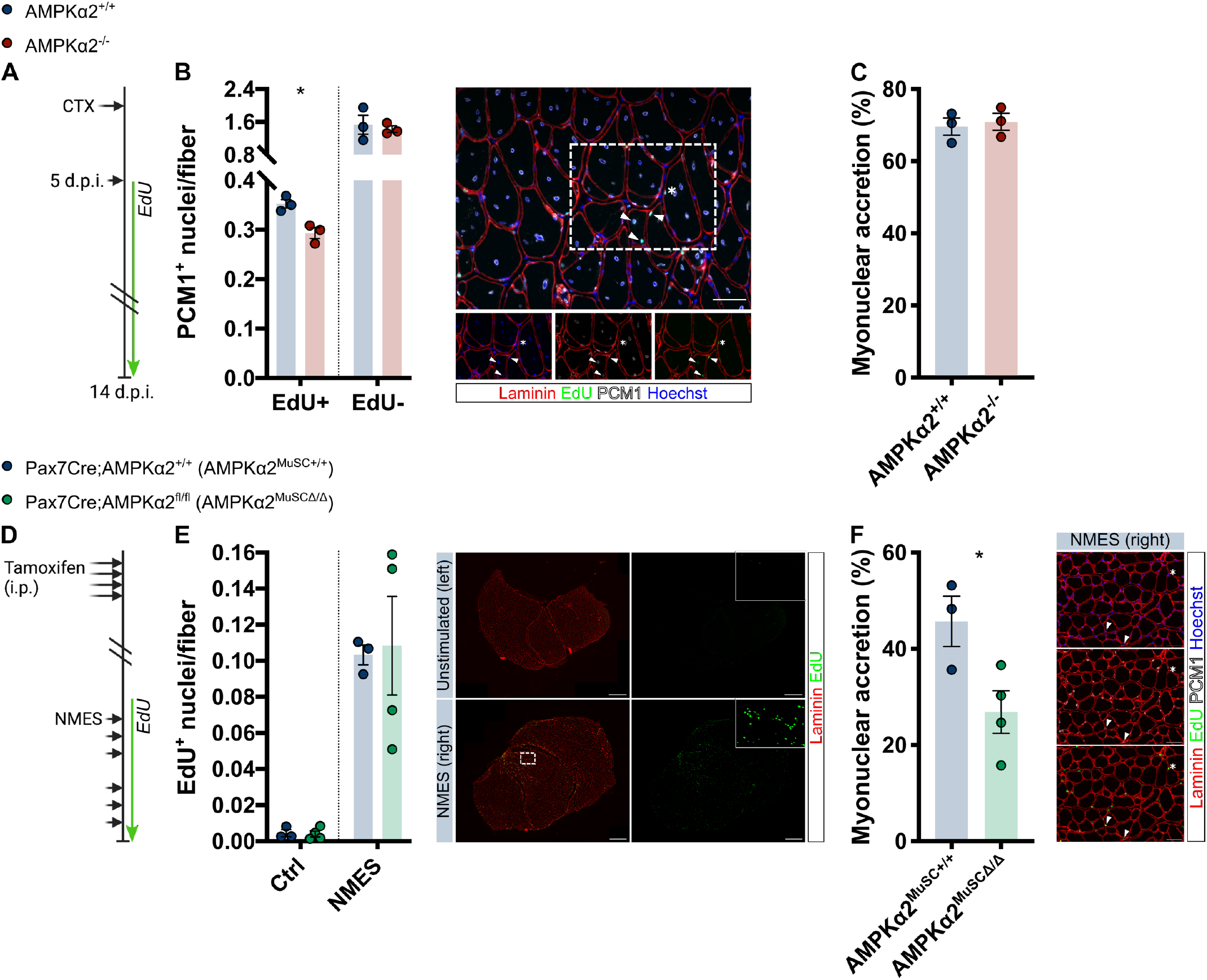
Role of AMPKα2 in the regulation of *in vivo* myonuclear accretion during regeneration and in response to muscle contraction induced by neuromuscular electrical stimulation (NMES). A. Schematic of EdU-based MuSC tracing in Cardiotoxin (CTX)-based skeletal muscle regeneration model in whole body AMPKα2^-/-^ mice. B. Stratification of Pericentriolar Material 1 (PCM1)^+^ nuclei per fiber by EdU staining at 14 days post injury (d.p.i.) (left panel), and representative image (right panel). Arrow heads indicate EdU^+^/PCM1^+^ nuclei, star indicates EdU^+^/PCM1^-^ nucleus. Length of white scale bar represents 50 µm. C. Percentage contribution of EdU^+^ nuclei below basal lamina to myonuclear accretion (*i*.*e*., are EdU^+^/PCM1^+^) at 14 d.p.i. D. Schematic of EdU-based MuSC tracing during NMES in mice after MuSC-specific AMPKα2 deletion. E. Number of EdU^+^ nuclei below the basal lamina in muscle subjected to NMES *versus* contralateral control muscle (Ctrl) (left panel), and representative images (right panel). Length of white scale bar represents 500 µm. White dashed square indicates area presented in Figure 4F. F. Percentage contribution of EdU^+^ nuclei below basal lamina to myonuclear accretion after NMES (left panel), and representative images (right panel). Arrow heads indicate EdU^+^/PCM1^+^ nuclei, star indicates EdU^+^/PCM1^-^ nucleus. Length of white scale bar represents 50 µm. Bars represent mean +/- SEM. Two-tailed, unpaired t-test. *p<0.05, compared to AMPKα2^+/+^ control. See also Figure S4.

To assess the MuSC *versus* myofiber-specific roles of AMPKα2 in the regulation of myonuclear accretion, we sought to trigger myonuclear accretion *in vivo* without injuring the myofibers. A recent study showed that 6 sessions of individualized NMES increased the number of nuclei per myofiber without invoking signs of myofiber damage or regeneration^42^. We resolved the MuSC *versus* myofiber-specific roles of AMPKα2 by subjecting cell-type specific conditional knockout mice to this individualized NMES protocol, and directly assessed myonuclear accretion by EdU-based cell tracing throughout the protocol (Figure 4D). Maximal *in vivo* tetanic force did not differ between wildtype and AMPKα2^MuSCΔ/Δ^ mice throughout the stimulation protocol (Figure S4F), imposing a similar relative stimulation load. Strikingly, after NMES, the number of EdU^+^ nuclei below the basal lamina (*i*.*e*., MuSCs & myofiber nuclei) were similarly increased in AMPKα2^MuSCΔ/Δ^ mice and controls (Figures 4E), indicating that MuSC activation and proliferation upon NMES are unaffected in absence of AMPKα2. However, their relative contribution to myonuclear accretion was substantially lower after AMPKα2 deletion from MuSCs (Figure 4F), resulting in a blunted increase in the number of PCM1^+^ nuclei after NMES (Figure S4G). Similar to AMPKα2^MuSCΔ/Δ^ mice, maximal *in vivo* tetanic force was unaltered in mice after myofiber-specific AMPKα2 deletion (AMPKα2^MFΔ/Δ^) throughout the stimulation protocol (Figures S4H and S4I). Furthermore, AMPKα2^MFΔ/Δ^ mice displayed a similar increase in EdU^+^ nuclei below the basal lamina compared to controls, as well as an unaffected myonuclear accretion percentage (Figures S4J and S4K), showing that AMPKα2 does not play an acute role in the myofiber-specific regulation of myonuclear accretion. Together, this shows a MuSC-intrinsic role of AMPKα2 in the regulation of myonuclear accretion upon myofiber contractions induced by NMES.

## DISCUSSION

MuSCs predominantly contribute to skeletal muscle homeostasis by two types of MuSC fusion; MuSC-MuSC fusion and myonuclear accretion. Myonuclear accretion is distinguished by the need for a coordination of cell type specific events between the myofiber and MuSCs, but it had remained unknown if there is a central molecular regulator that coordinates these events. Here, we demonstrate through *in vitro* and *in vivo* cell tracing that AMPKα2 specifically regulates myonuclear accretion in a MuSC-intrinsic manner.

Several previous studies proposed a specific regulation of myonuclear accretion^19–21,43–49^. However, a collective limitation is that they predominantly rely on *in vitro* data, and only one study (Noviello *et al*.^47^) directly assessed myonuclear accretion *in vivo*. Despite this limitation, published data collectively point towards a role for Ca^2+^ signalling in the distinct regulation of myonuclear accretion. Moreover, they unanimously show a regulation at the site of the myofiber, as Ca^2+^-dependent activation of CaMKII occurs in myotubes^21^, Ras Homolog Family Member A and SRF were shown to have a myofiber-intrinsic role^20,47^, and Nuclear factor of activated T-cells (NFAT) targets IL-4 and Stabilin-2 are specifically expressed by myotubes^43,48^. Thereby, they form a collective evidence that Ca^2+^ signalling contributes to myofiber-intrinsic regulation of myonuclear accretion.

In contrast to Ca^2+^-dependent activation of CaMKII^21^, AMPKα2 knockout did not alter gene expression of the fusogens *Mymk* and *Mymx*. Furthermore, AMPKα2 has not been reported to interact with mediators of these NFAT-related pathways. Moreover, in contrast to the myofiber-intrinsic regulation of myonuclear accretion observed in these studies, we observed no effect of myofiber-specific AMPKα2 knockout on either *in vitro* or *in vivo* myonuclear accretion. Rather, we showed a specific role of AMPKα2 in myocytes, which implies that AMPKα2-BAIAP2 signalling provides a distinct mechanism of regulation from NFAT-dependent pathways.

To our knowledge, only one study has previously indicated a myoblast-intrinsic specific regulation of myonuclear accretion, albeit *in vitro*. In this study, Teng *et al*., used cytoplasmic tracers to show that the formation of dual-labelled myotubes was impaired in co-cultures with Phospholipase D1 (PLD1) knockdown myoblasts, while PLD1 knockdown in myotubes had no effect^49^. Moreover, they showed that PLD1 knockout impairs the formation of myotubes with ≥5 nuclei, which can be partially rescued by treatment with lysophosphatidylcholine that facilitates opening and expansion of the fusion pore^49^. As there is evidence for reciprocal regulation between PLD1 and AMPK^50^, the AMPKα2-BAIAP2 axis described here may provide an alternative pathway of PLD1-mediated regulation of myonuclear accretion. In addition, regulation of PLD1 by AMPKα2 may provide a mechanism parallel to BAIAP2 by which AMPKα2 regulates myonuclear accretion.

Another target of AMPK, PGC1α, has been implicated as a regulator of myonuclear accretion upon endurance type exercise training^23^. However, exercise training in this study did not lead to an increase in the number of nuclei per fiber in wildtype mice, and an alteration in myonuclear accretion was extrapolated from a change in the correlation between the fiber CSA and number of nuclei per mm. Thus, while intriguing, a role of PCC1α as a regulator of myonuclear accretion – potentially controlled by AMPKα2 – should be confirmed by more direct measurements of this process.

In conclusion, our work reveals AMPKα2 as a novel MuSC-intrinsic regulator of myonuclear accretion. This adds to the cellular mechanisms by which AMPK controls the maintenance of skeletal muscle quality and quantity^22,51,52^. Blunted AMPK signalling is observed upon aging and in patients with metabolic disorders such as metabolic syndrome and COPD^53–55^, which are all conditions linked with impaired whole body metabolism and muscle pathology that may be improved upon AMPK activation. AMPK activation also mitigates pathologies of neuromuscular diseases such as Duchenne muscular dystrophy, spinal muscular atrophy, and myotonic dystrophy type 1^56^. Although technical limitations currently preclude the *in vivo* characterization of myonuclear accretion in these conditions, the identification of AMPKα2 as a regulator of myonuclear accretion may help to understand the pathophysiology and improve treatment of metabolic disorders and neuromuscular diseases.

### Limitations of (the) study

There is a paucity of studies directly assessing myonuclear accretion *in vitro* and *in vivo*, predominantly due to technical limitations. Several studies used cytoplasmic tracers to observe cell content mixing. However, this does not provide a direct measure of the number of nuclei donated to the myofiber (*i*.*e*., the rate of myonuclear accretion). Moreover, tracing of nuclei using nuclear fluorescent reporter proteins (*e*.*g*., H2B-GFP) proved unsuccessful due to nuclear propagation^57^. To overcome these limitations, we used EdU incorporation to trace MuSC-derived nuclei at the DNA level. However, EdU incorporation occurs during active DNA synthesis, and as such, we only trace MuSCs that undergo cell division. Although it is unknown if and to what extend MuSCs fuse without prior cell division, it may lead to an underestimation of the effects observed here.

Given the technical difficulties in measuring myonuclear accretion, and the fact that myofibers are the distinguishing fusion partners in this process, the majority of the identified specific regulators of myonuclear accretion to date act in a myofiber-intrinsic manner. Using specific *in vitro* and *in vivo* myonuclear accretion assays, we showed that AMPKα2-BAIAP2 signalling specifically regulates myonuclear accretion in a MuSC-intrinsic manner. Yet, the physiological purpose of MuSC-intrinsic regulation of myonuclear accretion remains unknown. We could speculate that such regulators promote processes such as actin cytoskeletal rearrangements that may be unique to myonuclear accretion, as suggested in drosophila^58^. Alternatively, such regulation may be part of a cellular cross-talk, where myofibers induce distinct signalling in MuSCs to prioritize nuclear recruitment over the establishment of new myofibers. This would be in line with the two successive waves of differentiation identified during postnatal muscle growth^59^, and should be a topic of further investigation.

## Supporting information

Video S1 fusion after differentiation

Video S2 motility after differentiation

Video S3.1 myonuclear accretion in coculture +991

Video S3.2 myonuclear accretion in coculture +991

Video S3.3 myonuclear accretion in coculture +991

Video S3.4 myonuclear accretion in coculture +991

## ACKNOWLEDGEMENTS

This work was supported by Centre National de la Recherche Scientifique, the Association Française Contre les Myopathies-Téléthon (Alliance MyoNeurALP1 and MyoNeurALP2) and the Agence Nationale de la Recherche (ANR-22-CE14-0032-01). AK was supported by a Kootstra Talent Fellowship (Maastricht University) and a Marie Skłodowska-Curie Individual Fellowship (#896544). AS was supported by Fondation pour la Recherche Médicale (#ECO202206015552). LG was supported by Ligue Contre le Cancer (#GB/MA/SC 12627)

## AUTHOR CONTRIBUTIONS

RM designed the study. AK and RM conceived, performed and analyzed experiments. MT, SBL, LG, AS, CD and AF performed and analyzed experiments. AK prepared the figures and wrote the manuscript. RM, MT and KS provided conceptual input and edited the manuscript. All authors read and approved the manuscript.

## METHODS

### Mice

Mice were bred, housed and maintained in accordance with the French and European legislation. The experimental protocols were approved by the local ethical committee. Experiments were conducted on males at 8-12 weeks of age. Mice were genotyped by PCR using toe or tail DNA. AMPKα2 mice were used for MuSC extraction and *in vivo* experiments^60^. Pax7Cre;AMPKα2^fl/fl^ were obtained by crossing Pax7CreER^T2/+^ mice^61^ with AMPKα2^fl/fl^ mice^60^. Similarly HSACre;AMPKα2^fl/fl^ were obtained by crossing HSACreER^T2/+^ mice^62^ with AMPKα2^fl/fl^ mice. Activation of Cre activity in CreER^T2^ mice was induced by daily intraperitoneal (i.p.) Tamoxifen (MP Biochemicals #0215673891; 0.1mg/g BW in sunflower oil) injections for 4 days. Subsequent experiments were initiated 1 week (*i*.*e*., for regeneration experiment) or 3 weeks (*i*.*e*., for NMES) after the first Tamoxifen injection. Skeletal muscle injury was induced by Cardiotoxin (CTX; Latoxan #L8102) injection in the *Tibialis Anterior* (TA) (50µl per TA, 12µM). Mice were fed 5 ethynyl 2’ deoxyuridine (EdU) in the drinking water (Carbosynth #NE0870; 0.5 mg/ml, 1% glucose) at indicated timepoints and durations to label and trace activated MuSCs.

### Muscle force measurement and NMES

*In situ* muscle force production was assessed as described previously^63^. Briefly, mice were anaesthetized by i.p. pentobarbital sodium injection (Ref; 50 mg/kg). The knee and foot were fixed with clamps and stainless-steel pins, and the distal tendon of the TA was cut and attached to an isometric transducer (Harvard Bioscience) using a silk ligature. The sciatic nerves were proximally crushed and distally stimulated by bipolar silver electrode using supramaximal square wave pulses of 0.1 ms duration. Maximal force production was determined in response to isometric contractions induced by 500 ms stimulation trains at 75 150 Hz. To assess fatigue resistance, muscles were stimulated at 20 Hz. Fatigue resistance was expressed as time to reach <70% of initial force.

Myonuclear accretion was invoked by *in vivo* individualized non-damaging neuromuscular electrical stimulation (NMES) of the mouse plantar flexor muscles under isoflurane aneasthesia^42^. Briefly, mice were subjected to 2*3 consecutive days of NMES consisting of 80 stimulation trains at 50 Hz (2 s duration, 8 s recovery) at a current intensity resulting in 15% of maximal isometric force production (Fmax). Fmax was measured in response to a 250 ms stimulation train at 100 Hz. Mice were sacrificed after an Fmax test, 24 hr after the last NMES session.

### MuSC isolation

MuSCs were isolated from hindlimb muscles of AMPKα2^+/+^ and AMPKα2^-/-^ mice, using an adaptation of published methods^26,64^. Briefly, muscles were dissected and digested with Collagenase II (Gibco #17101015) and Dispase (Gibco #17105041)^64^. Dissociated muscles were passed through a 70µm filter and red blood cells were lysed in ACK lysis buffer (Lonza #10 548E). Cell suspensions were then incubated with anti CD45, anti CD31, anti Sca1, anti CD34 (all eBioscience), and anti α7integrin (AbLab), and CD45/CD31/Sca1^-^;CD34/α7int^+^ were sorted using a BD FACSAria II cytometer (FACS)^26^.

Alternatively, cell suspensions were incubated with Satellite Cell Isolation Kit (Miltenyi Biotec #130 104 268) for non MuSC depletion by magnetic activated cell sorting (MACS). MuSC purity after MACS was verified by Desmin IHC to be at 94±3% (mean±SD).

### MuSC primary culture

FACS isolated MuSCs were seeded onto Matrigel (Corning, #354234) coated supports at 3000 cells/cm^2^, and amplified during 5 days in proliferation medium (DMEM F12 (Gibco #31331028), 20% Foetal Bovine Serum (FBS) (Gibco #10270106), 2% Ultroser G (PALL life sciences #15950017), and 1% Penicillin/Streptomycin (PS) (Gibco #15140122). Proliferation medium was replaced at day 3 of amplification.

MACS isolated MuSCs were seeded onto Gelatin (Sigma #G1393) coated supports at 3000 cells/cm^2^, and amplified ≤3 passages in proliferation medium. Only experiments presented in Figure 3G were conducted using MACS-isolated MuSCs.

After amplification, MuSC progeny (*i*.*e*., myoblasts) were trypsinized and plated at indicated densities onto Matrigel coated supports. To induce differentiation, proliferation medium was removed and replaced by differentiation medium (DMEM F12, 2% Horse serum (HS) (Gibco #16050130), and 1 % PS). To trace myoblasts *in vitro*, cells were incubated with EdU (Invitrogen #C10350; 10 µM). To induce AMPK activation, cells were treated with the small molecule compound 991 (Selleckchem #S8654; 10 µM). Myoblasts were transfected with pECE M2 BAIAP2 WT (BAIAP2 WT) or pECE M2 BAIAP2 S366A (BAIAP2^S366A^) using Lipofectamine 2000 (Invitrogen #11668019). 6 hr after transfection, cells were induced to differentiate. Plasmids were a gift from Anne Brunet (addgene #31656, http://n2t.net/addgene:31656, RRID:Addgene_31656; addgene # 31657, http://n2t.net/addgene:31657, RRID:Addgene_31657).

### Immunohistochemistry

For immunohistochemical analyses, muscles of interest were isolated, embedded in tragacanth gum (VWR #ICNA0210479280), frozen in liquid nitrogen cooled isopentane, and stored at -80°C until use. 10 µm-thick cryosections were prepared for hematoxylin-eosin (HE) staining and immunolabeling. HE staining was used to assess the efficiency of CTX injections, and muscles were only kept for further analyses if ≥75% of the muscle area consisted of centrally nucleated fibers. For immunolabeling, cryosections were permeabilized for 10 min in 0.5% Triton X-100 in PBS and saturated in 2% BSA for 1 hr at room temperature (RT). For identification of regenerating myofibers, sections were co-labelled overnight at 4°C with primary antibodies directed against embryonic Myosin Heavy Chain (eMHC; Santa Cruz #sc-53091; 1:100) and Laminin α1 (Sigma-Aldrich #L9393; 1:200). For identification of myofiber nuclei, sections were co-labelled with primary antibodies directed against Pericentriolar Material 1 (PCM1; Sigma-Aldrich #HPA023370; 1:1000) and Laminin α2 (Santa Cruz #sc-59854; 1:1000). For identification of MuSCs, sections were fixed for 20 min in 4% paraformaldehyde (PFA) at RT, and permeabilized for 6 min in 100% methanol at -20°C. After antigen retrieval for 2*5 min in 10 mM Citrate buffer at 90°C, sections were saturated in 4% BSA for 2 hr at RT. Sections were then co-labelled overnight at 4°C with primary antibodies directed against Paired Box 7 (Pax7; DSHB; 1:50) and Laminin α1 (Sigma-Aldrich #L9393; 1:100) in 2% BSA. Secondary antibodies were coupled to FITC, Cy3 or Cy5 (Jackson ImmunoResearch Inc.; 1:200) and incubated for 45-60 min at 37°C. Labelling of EdU was performed after secondary antibody incubation using Click-It EdU Kit (ThermoFisher Scientific #c10337). Nuclear counterstain was performed by 10 sec incubation with Hoechst (Sigma-Aldrich #14533; 2 µM), and coverslips were mounted with Fluoromount G (Interchim, #FP-483331).

Cultured cells were fixed for 10 min in 4% PFA at RT, permeabilized for 10 min in 0.5% Triton X-100 in PBS, and saturated in 4% BSA for 1 hr at room temperature (RT). Cells were then incubated overnight at 4°C with primary antibodies directed against Myogenin (MyoG; Santa Cruz #SC-12732; 1:50) or Desmin (Abcam #Ab32362; 1:200) in 2% BSA. Secondary antibodies were coupled to FITC, Cy3 or Cy5 (Jackson ImmunoResearch Inc.; 1:200) and incubated for 45-60 min at 37°C. Labelling of EdU was performed after secondary antibody incubation using Click-It EdU Kit (ThermoFisher Scientific #c10337). Nuclear counterstain was performed by 10 sec incubation with Hoechst (Sigma-Aldrich #14533; 2 µM), and cells were covered with Fluoromount G (Interchim, #FP-483331).

Images of fluorescent immunolabeling in cell culture supports and scanned slides were acquired with a Zeiss Axio Observer Z1 connected to a Coolsnap HQ2 camera. Scanned slides of PCM1 stainings were acquired with a Zeiss Axio Scan.Z1. Randomly chosen fields of view from sections or cells cultured on removable chamber slides were acquired using a Zeiss Axio Imager Z1 connected to a Coolsnap Myo camera. For each condition of each experiment, at least 5-10 randomly chosen fields of view were counted. The fusion index was calculated as the percentage of nuclei within a cell containing ≥2 nuclei. *In vitro* myonuclear accretion was defined as fusion between an EdU-labelled myocyte and a pre-generated myotube (≥2 nuclei), and was calculated as the percentage of EdU^+^ nuclei within a myotube containing ≥2 EdU^-^ nuclei.

### Live cell imaging

Time-lapse imaging of live cells was performed using a Zeiss Axio Observer Z1 connected to a Coolsnap HQ2 camera. For analysis of cell motility, myocytes were captured by brightfield imaging at 5 min intervals. At least 50 cells were tracked in 5-10 individual fields of view covering a total of 3 hr using MetaMorph image analysis software. For analysis of fusion events, cells were captured by brightfield imaging at 15 min intervals. Fusion events of at least 50 myocytes were scored in 5 individual fields of view covering a total of 12 hr. To observe myonuclear accretion, myocytes were labelled with 1 µM CellTracker Deep Red (ThermoFisher #C34565) according to the manufacturer instructions, and then co-cultured with 24-hour differentiated unstained myotubes. 12 hr after initiation of co-culture, cells were captured by brightfield and fluorescent imaging at 6.7 min intervals covering a total of 5 hr.

### AMPK activity

Skeletal muscle tissue (*Tibialis Anterior*) was rapidly harvested from the indicated animals, snap-frozen in liquid nitrogen, and stored at -80°C. The muscles were homogenized in cold lysis buffer, and protein extracts were prepared as previously described^28^. AMPKα2 *versus* AMPKα1-containing complexes were immunoprecipitated using an anti-AMPKα2 and anti-AMPKα1 antibody, respectively, and *in vitro* phosphotransferase activity was determined towards the AMRA peptide (AMRAASAAALARRR) as previously described^28^.

### Verification of AMPKα2 deletion by PCR

To verify MuSC specific deletion of AMPKα2, hindlimb muscles of Tamoxifen treated Pax7Cre;AMPKα2^fl/fl^ mice were collected 3 weeks after the first Tamoxifen injection. MuSCs were extracted by FACS sorting of CD45/CD31/Sca1^-^;CD34/α7int^+^ cells, and CD45/CD31/Sca1^+^ were sorted as non MuSC controls. For HSACre;AMPKα2^fl/fl^ mice, deletion was verified at the end of the experimental protocol in whole muscle. DNA was extracted in 25 mM NaOH, 0.2 mM EDTA, and neutralized in 40 mM Tris-HCL ph5. Amplification was performed using REDExtract N Amp PCR ReadyMix (Sigma #R4775) according to the manufacturer protocol.

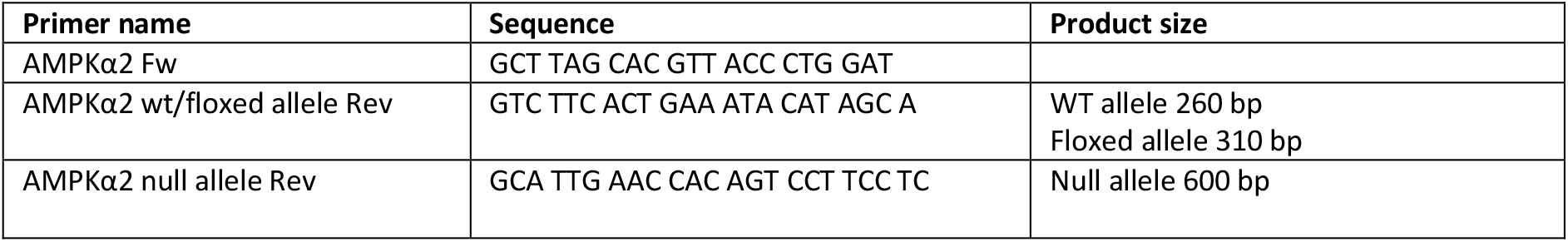

### RT qPCR

Cells were lysed in TRIzol reagent (Invitrogen #15596026), and total RNA was obtained by chloroform/isopropanol based extraction according to the manufacturer protocol. RNA was reverse transcribed using Superscript II Reverse Transcriptase (Invitrogen #18064022), and qPCR was carried out using 1.5 µl of cDNA, 5 µl LightCycler 480 SYBR Green I Master (Roche #04887352001), and 0.5 µl primers (10 µM) at 10 µl total volume. After initial 5 min denaturation at 95°C, amplification was performed at 95°C (10 sec), 60°C (5 sec), 72°C (10 sec) for 45 cycles carried out on a Bio Rad CFX. Relative gene expression was calculated using 2 ^ΔΔCt^. Values were normalized to the geometric mean of tree reference genes (PPI, RPLP0, and B2M).

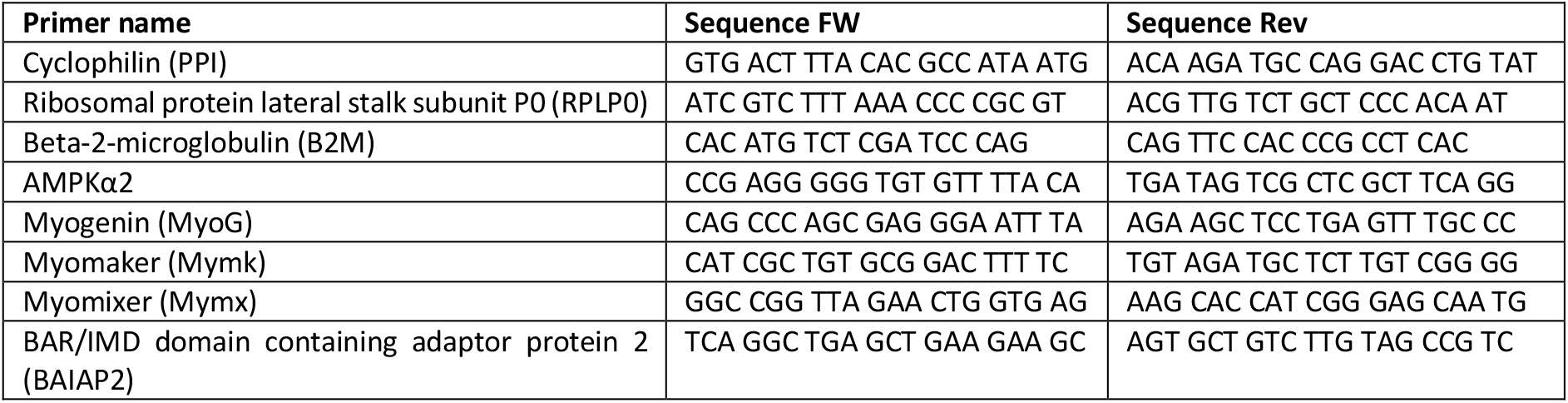

### Statistical analyses

For *in vitro* experiments, replicates signify independent experiments. For *in vivo* experiments, replicates signify individual muscles. All *in vivo* results were analysed using unpaired parametric analyses, whereas *in vitro* results were analysed using paired parametric analyses. Statistical significance was determined using two-sided Student’t *t* tests or ANOVA with Bonferroni post hoc analyses in case of multiple comparisons. Time points were compared if specifically indicated.

---

**Figure S1.**
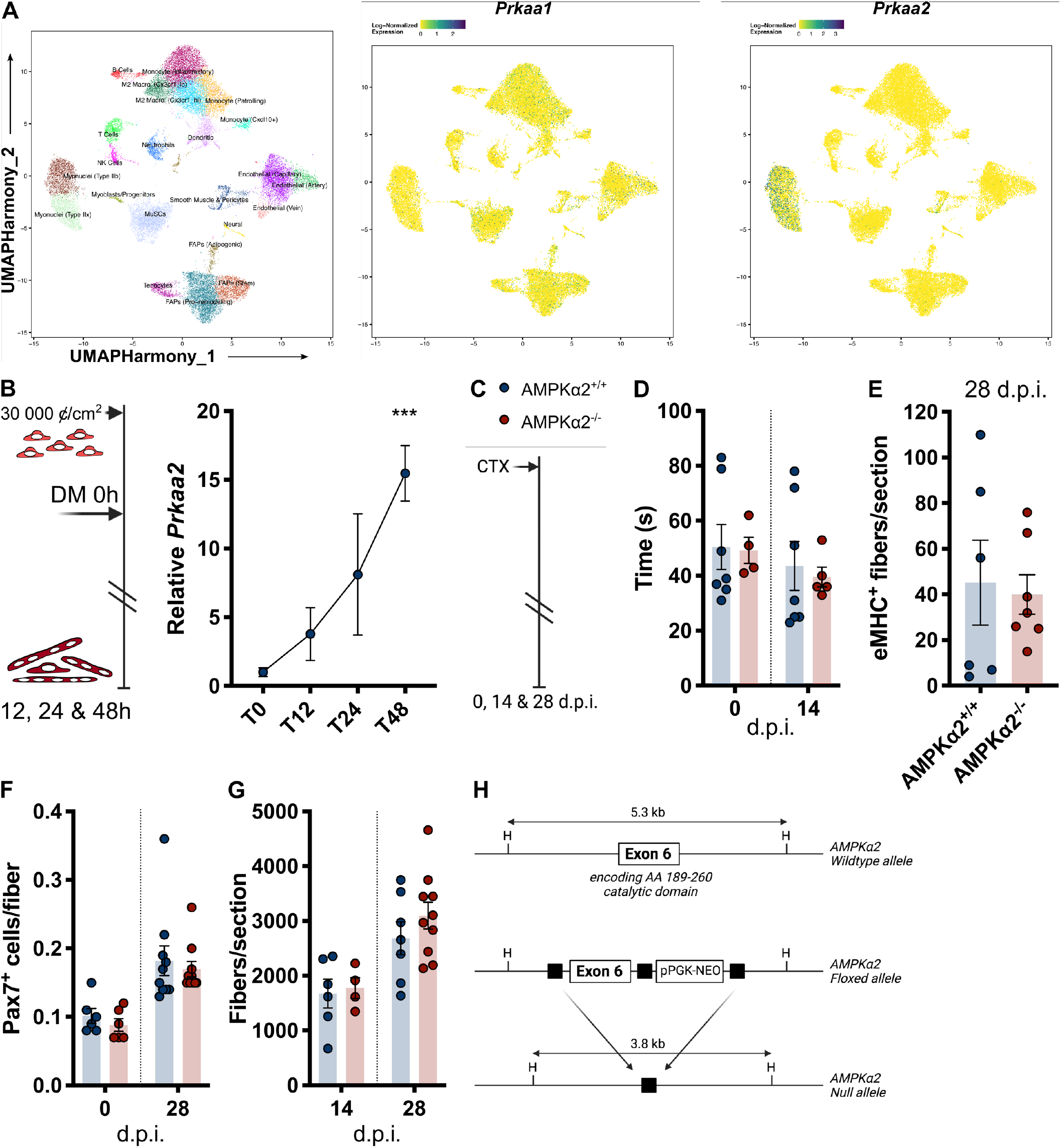
Role of AMPKα2 in skeletal muscle regeneration. A. Harmony integrated UMAP of single cell/single nucleus RNA sequencing (scMuscle version 1.1)^25^, displaying cell populations (left panel), expression of *Prkaa1* (middle panel), and expression of *Prkaa2* (right panel). Skeletal muscle regeneration in whole body AMPKα2^-/-^ mice (B-F). B. Schematic of CTX-based skeletal muscle regeneration model and analysis endpoints at indicated d.p.i. C. Time (seconds) until force production <70% of initial force before and 14 d.p.i. D. Number of eMHC stained fibers per section at 28 d.p.i. E. Number of Pax7 stained cells per fiber before and 28 d.p.i. F. Number of fibers per section at 14 and 28 d.p.i. G. Relative *Prkaa2* gene expression during *in vitro* myogenesis. Left panel shows schematic representation of experiment. H. Schematic of AMPKα2 gene targeting. Black boxes indicate LoxP sites, H’s indicate HindIII restriction sites. Bars represent mean +/- SEM. In line chart, dots represent mean +/- SEM. Two-tailed, paired t-test, ***p<0.001, compared to 0 d.p.i.

**Figure S2.**
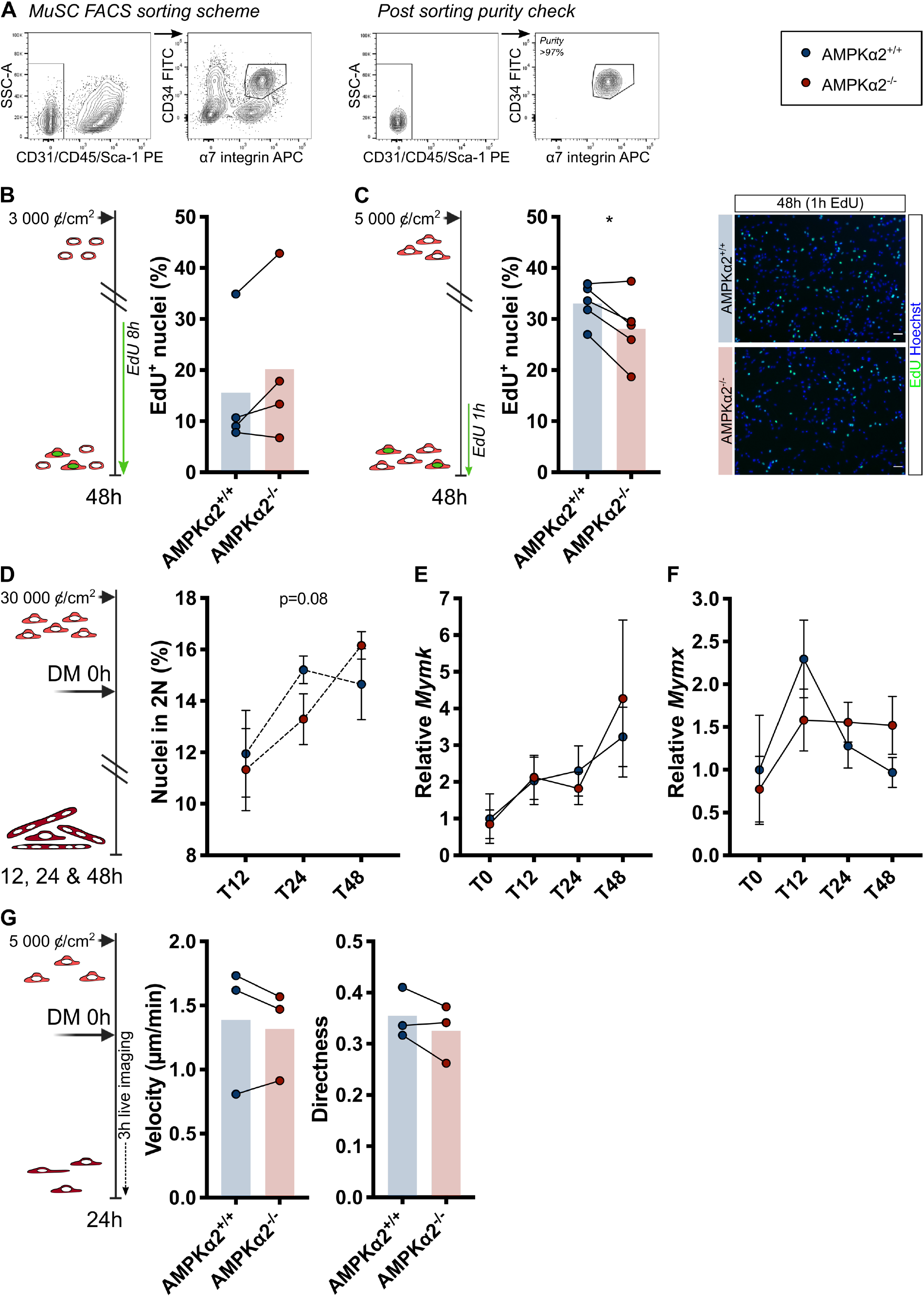
Role of AMPKα2 in *in vitro* myogenesis. Left panels show schematic representation of experiments. A. Schematic of gating for MuSC sorting by FACS (left panels). MuSCs population (CD45/CD31/Sca1^-^;CD34/α7int^+^) is at >97% purity after sorting (right panels). B. Percentage of EdU stained nuclei during MuSC activation. C. Percentage of EdU stained nuclei during myoblast proliferation (middle panel), and representative images (right panel). D. Percentage of nuclei in cells with 2 nuclei during *in vitro* myogenesis. E. Relative *Myomaker* (*Mymk)* gene expression during *in vitro* myogenesis. F. Relative *Myomixer* (*Mymx)* gene expression during *in vitro* myogenesis. G. Cell motility of myoblasts during differentiation in non-fusing cultures expressed as cell velocity (middle panel) and directness (*i*.*e*., Euclidian distance/total distance) (right panel). Bars represent mean. In line chart, dots represent mean +/- SEM. Two-tailed, paired t-test, *p<0.05, compared to AMPKα2^+/+^ control at same timepoint. Length of white scale bars represent 50 µm.

**Figure S3.**
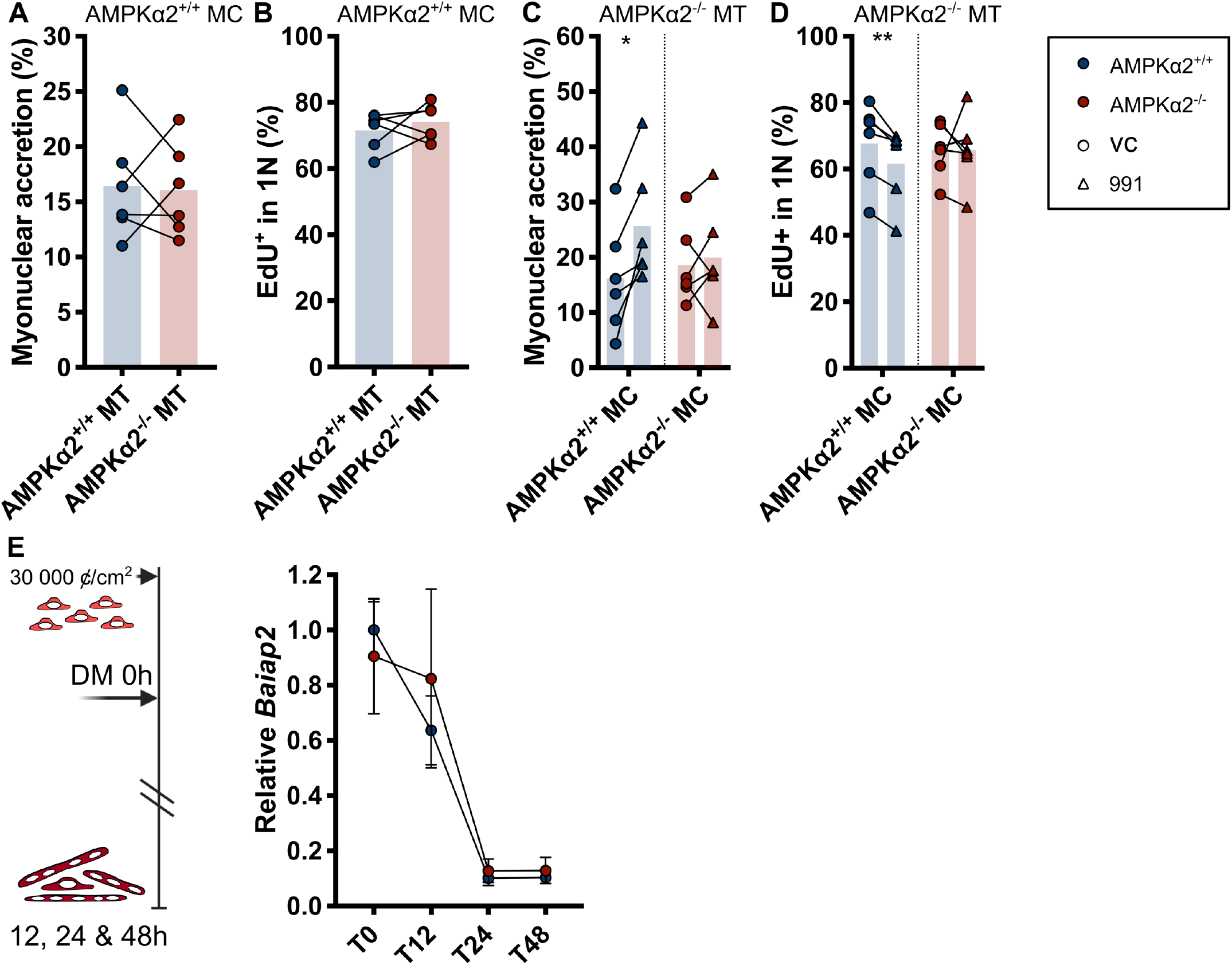
Myotube-intrinsic regulation of *in vitro* myonuclear accretion by AMPKα2. Co-cultures between AMPKα2^+/+^ *versus* AMPKα2^-/-^ myotubes and AMPKα2^+/+^ myocytes (A-B). MC=myocytes, MT=myotubes. A. Percentage contribution of EdU^+^ myocytes to myonuclear accretion after 48 hours of co-culturing. B. Percentage EdU^+^ myocytes that remain mononucleated after 48 hours of co-culturing. Co-cultures between AMPKα2^-/-^ myotubes and AMPKα2^+/+^ *versus* AMPKα2^-/-^ myocytes (C-D). C. Percentage contribution of EdU^+^ myocytes to myonuclear accretion after 48 hours of co-culturing with 10 µM 991 or vehicle control (VC). D. Percentage EdU^+^ myocytes that remain mononucleated after 48 hours of co-culturing with 10 µM 991 or vehicle control (VC). E. Relative *Baiap2* gene expression during *in vitro* myogenesis of myoblasts derived from MuSCs isolated by MACS. Bars represent mean. In line chart, dots represent mean +/- SEM. Two-tailed, paired t-test, *p<0.05, **p<0.01, compared to VC.

**Figure S4.**
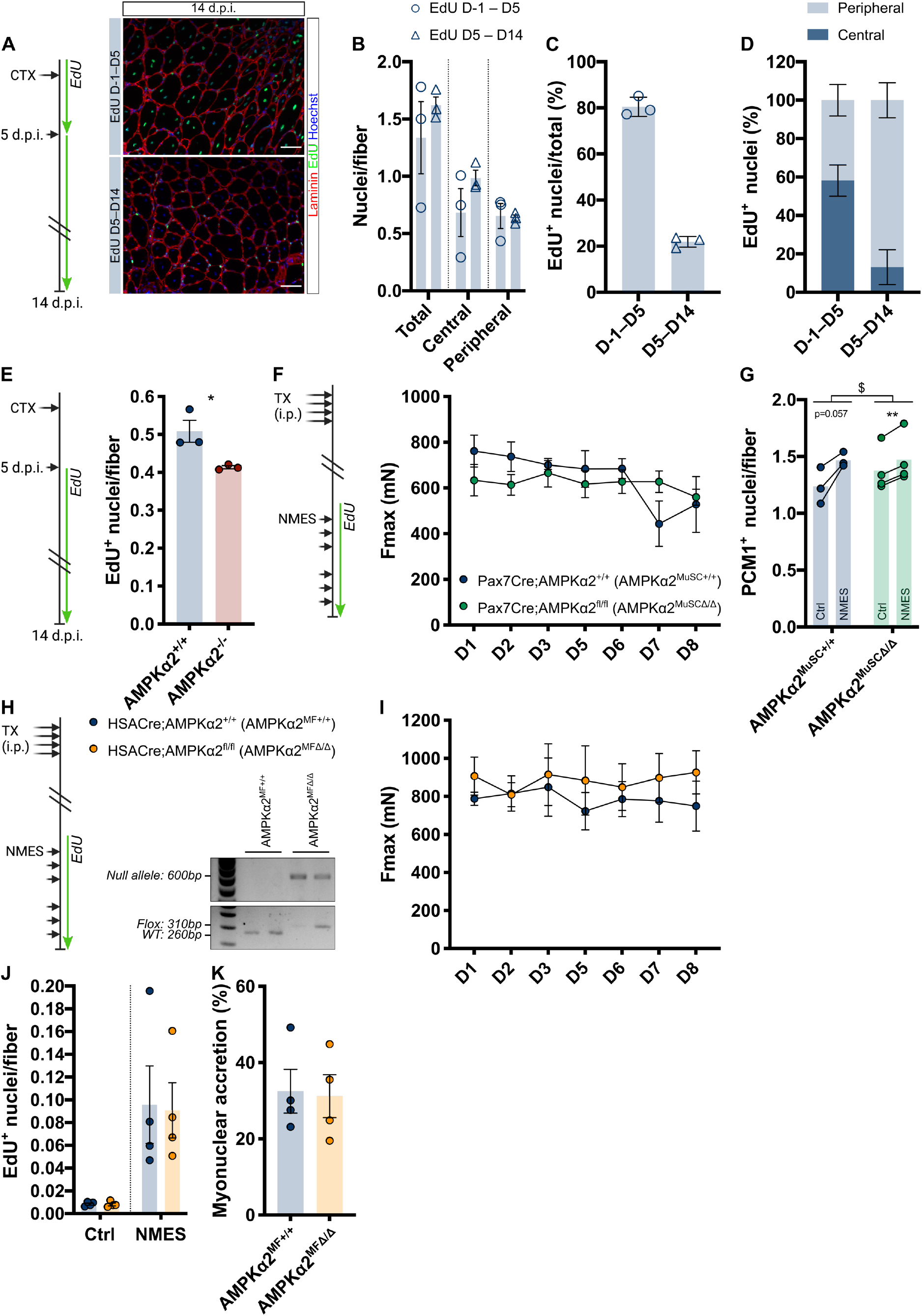
Role of AMPKα2 in the regulation of *in vivo* myonuclear accretion during regeneration and in response to NMES. A. Schematic of EdU-based MuSC tracing in CTX-based skeletal muscle regeneration model during day -1–5 *versus* 5-14 d.p.i. in wildtype mice (left panel), and representative images (right panel). B. Number and localization of nuclei below the basal lamina at 14 d.p.i. C. Percentage of nuclei below the basal lamina that are EdU^+^ at 14 d.p.i. D. Relative localization of EdU^+^ below the basal lamina at 14 d.p.i. E. Number of EdU^+^ nuclei below the basal lamina at 14 d.p.i. in AMPKα2^+/+^ and AMPKα2^-/-^ mice. F. Maximal plantar flexor muscle force at indicated NMES sessions in mice after MuSC-specific AMPKα2 deletion. G. Number of PCM1^+^ nuclei per fiber in muscle subjected to NMES (NMES) *versus* contralateral control muscle (Ctrl). H. Verification of recombination in whole muscle of HSACre;AMPKα2^+/+^ (AMPKα2^+/+^) and HSACre;AMPKα2^fl/fl^ (AMPKα2^Δ/Δ^) mice after NMES. I. Maximal plantar flexor muscle force at indicated NMES sessions in mice after myofiber-specific AMPKα2 deletion. J. Number of EdU^+^ nuclei below the basal lamina in muscle subjected to NMES *versus* contralateral control muscle (Ctrl). K. Percentage contribution of EdU^+^ nuclei below basal lamina to myonuclear accretion after NMES. Bars represent mean +/- SEM. In line chart, dots represent mean +/- SEM. Two-way ANOVA; ^$^p<0.05 genotype*NMES effect. Two-tailed, paired t-test, **p<0.01, compared to contralateral Ctrl. Two-tailed, unpaired t-test; *p<0.05, compared to AMPKα2^+/+^ control. Length of white scale bars represent 50 µm.

**Video S1**.

Fusion of myocytes during 12 hours of live imaging.

**Video S2**.

Cell motility of myoblasts during differentiation in non-fusing cultures.

**Video S3.1-3.4**.

*In vitro* myonuclear accretion in co-cultures between myocytes and myotubes treated with the AMPK activator 991.

